# Genome-wide meta-analysis of depression identifies 102 independent variants and highlights the importance of the prefrontal brain regions

**DOI:** 10.1101/433367

**Authors:** David M. Howard, Mark J. Adams, Toni-Kim Clarke, Jonathan D. Hafferty, Jude Gibson, Masoud Shirali, Jonathan R. I. Coleman, Saskia P. Hagenaars, Joey Ward, Eleanor M. Wigmore, Clara Alloza, Xueyi Shen, Miruna C. Barbu, Eileen Y. Xu, Heather C. Whalley, Riccardo E. Marioni, David J. Porteous, Gail Davies, Ian J. Deary, Gibran Hemani, Klaus Berger, Henning Teismann, Rajesh Rawal, Volker Arolt, Bernhard T. Baune, Udo Dannlowski, Katharina Domschke, Chao Tian, David A. Hinds, 23andMe Research Team, Major Depressive Disorder Working Group of the Psychiatric Genomics Consortium, Maciej Trzaskowski, Enda M. Byrne, Stephan Ripke, Daniel J. Smith, Patrick F. Sullivan, Naomi R. Wray, Gerome Breen, Cathryn M. Lewis, Andrew M. McIntosh

## Abstract

Major depression is a debilitating psychiatric illness that is typically associated with low mood, anhedonia and a range of comorbidities. Depression has a heritable component that has remained difficult to elucidate with current sample sizes due to the polygenic nature of the disorder. To maximise sample size, we meta-analysed data on 807,553 individuals (246,363 cases and 561,190 controls) from the three largest genome-wide association studies of depression. We identified 102 independent variants, 269 genes, and 15 gene-sets associated with depression, including both genes and gene-pathways associated with synaptic structure and neurotransmission. Further evidence of the importance of prefrontal brain regions in depression was provided by an enrichment analysis. In an independent replication sample of 1,306,354 individuals (414,055 cases and 892,299 controls), 87 of the 102 associated variants were significant following multiple testing correction. Based on the putative genes associated with depression this work also highlights several potential drug repositioning opportunities. These findings advance our understanding of the complex genetic architecture of depression and provide several future avenues for understanding aetiology and developing new treatment approaches.

---

Depression is the leading cause of worldwide disability ^1^ and an estimated 1 in 6 people will develop the disorder during their lifetime ^2^. Twin studies have provided heritability estimates of the disease of approximately 30-40% ^3^, however depression is a polygenic trait influenced by many genetic variants each of small effect ^4^. Therefore, to enable the detection of causal genetic variants associated with depression there is a need to study very large numbers of individuals. However, obtaining detailed clinical diagnoses of major depressive disorder in larger cohorts is both time consuming and expensive. The results of Howard, et al. ^5^ showed that there is a strong genetic correlation (r_G_ = 0.86, s.e. = 0.05) between broader self-declared definitions of depression and clinically diagnosed major depressive disorder (MDD) within a hospital setting. Therefore, analysing larger samples, which have used different approaches to diagnosis, may provide advances in our understanding of the genetics of depression.

Major efforts to identify genetic variants associated with depression have included a mega-analysis of 9 cohorts (total n = 18,759; 9240 cases and 9519 controls) for MDD ^4^ and a meta-analysis of 17 cohorts (total n = 34,549) using a broader diagnostic scale that includes depressive symptoms ^6^. However, both of these studies failed to find any replicated variants associated with depression. The first study to report replicable genetic variants for depression found two significant loci associated with severe, recurrent MDD (85% enriched for melancholia) in a sample of Han Chinese women (total n = 10,640; 5,303 cases and 5,337 controls) ^7^. A later study conducted by Hyde, et al. ^8^, examining research participants from the personal genetics company 23andMe, Inc., used a self-reported clinical diagnosis of depression as the phenotype and identified 15 associated loci (total n = 459,481; 121,380 cases and 338,101 controls). More recently, a genome-wide association analysis of UK Biobank by Howard, et al. ^5^ identified 17 variants associated across three depression phenotypes (maximum total n = 322,580; 113,769 cases and 208,811 controls). These three depression phenotypes ranged from self-reported help-seeking for problems with nerves, anxiety, tension or depression (termed “broad depression”), probable MDD based on self-reported depressive symptoms with associated impairment, and MDD identified from hospital admission records. Finally, a meta-analysis of 35 cohorts (total n = 461,134; 130,664 cases and 330,470 controls) conducted by Wray, et al. ^9^ (PGC) found 44 loci that were significantly associated with a spectrum of depression phenotypes, some obtained from structured clinical interview and others based on broader criteria.

To maximise the power of the present study we conducted a genome-wide meta-analysis of depression using 807,553 individuals (246,363 cases and 561,190 controls, after excluding overlapping samples) from the three largest studies noted above ^5, 8, 9^. From the Hyde, et al. ^8^ analysis, our meta-analysis included only the 23andMe discovery cohort (termed “23andMe_307k”; 75,607 cases and 231,747 controls). From the Howard, et al. ^5^ analysis of UK Biobank, the “broad depression” phenotype was included, with 4-means clustering of genomic principal components used to derive a larger sample (127,552 cases and 233,763 controls) than studied previously. The PGC analysis ^9^ included the 23andMe_307k discovery cohort and an earlier data release of the UK Biobank cohort (n = 29,740, 14,260 cases and 15480 controls); we therefore obtained results from the PGC that excluded both of these cohorts (termed “PGC_139k”; 43,204 cases and 95,680 controls).

We sought replication of the variants associated with depression within a set of 23andMe participants independent of the 23andMe_307k cohort included in the meta-analysis (414,055 cases of self-reported clinical diagnosis of depression and 892,299 controls). The results from the meta-analysis were used to calculate genetic correlations and conduct Mendelian randomization to identify potentially pleiotropic and causal relationships between depression and other diseases and behavioural traits. The meta-analysis results were also used to identify a set of associated genes and gene-pathways, as well as enrichment of functional annotations associated with depression. Combining evidence of enrichment in biological pathways with information on traits correlated with depression allows for additional inferences about shared aetiological mechanisms, thereby increasing the utility of the standard association analysis approach. Interactions between associated genes and available drug treatments were also examined to identify novel drug treatments for depression.

## Results

### 102 independent genetic variants associated with depression

We conducted a genome-wide association meta-analysis of depression using 807,553 individuals (Table 1; 246,363 cases and 561,190 controls) from three previous studies of depression, after removing sample overlap; the previous studies were Hyde, et al. ^8^ Howard, et al. 5, and Wray, et al. ^9^. We tested the effects of 8,098,588 genetic variants on depression and identified 9,744 associated variants (*P* < 5 × 10^-8^) of which 102 variants in 101 loci were independently segregating (Supplementary Table 1). The basepair positions of these loci were identified by clumping all associated variants (linkage disequilibrium r^2^ < 0.1 across a 3 Mb window) and then merging any overlapping clumps. Independent variants in each locus were identified through conditional analysis ^10^ using all variants in that locus.

**Table 1.** Sample sizes and the proportion of males and females of the depression cohorts used in the meta-analysis and the replication cohort

| **Cohort** | **Study** | **Cases**<br>**(Prop. Male, Female)** | **Controls**<br>**(Prop. Male, Female)** | **Total**<br>**(Prop. Male, Female)** |
| --- | --- | --- | --- | --- |
| 23andMe_307k | Hyde, et al. <sup>8</sup> | 75,607 (0.38, 0.62) | 231,747 (0.56, 0.44) | 307,354 (0.52, 0.48) |
| UK Biobank | Howard, et al. <sup>5</sup> | 127,552 (0.35, 0.65) | 233,763 (0.52, 0.48) | 361,315 (0.46, 0.54) |
| PGC_139k† | Wray, et al. <sup>9</sup> | 43,204 | 95,680 | 138,884 |
| **Meta-analysis** |  | **246,363** | **561,190** | **807,553** |
| Replication | Unpublished | 414,055 (0.30, 0.70) | 892,299 (0.50, 0.50) | 1,306,354 (0.44, 0.56) |

† Proportion of males and females not reported for PGC\_139k

**Table 2.**
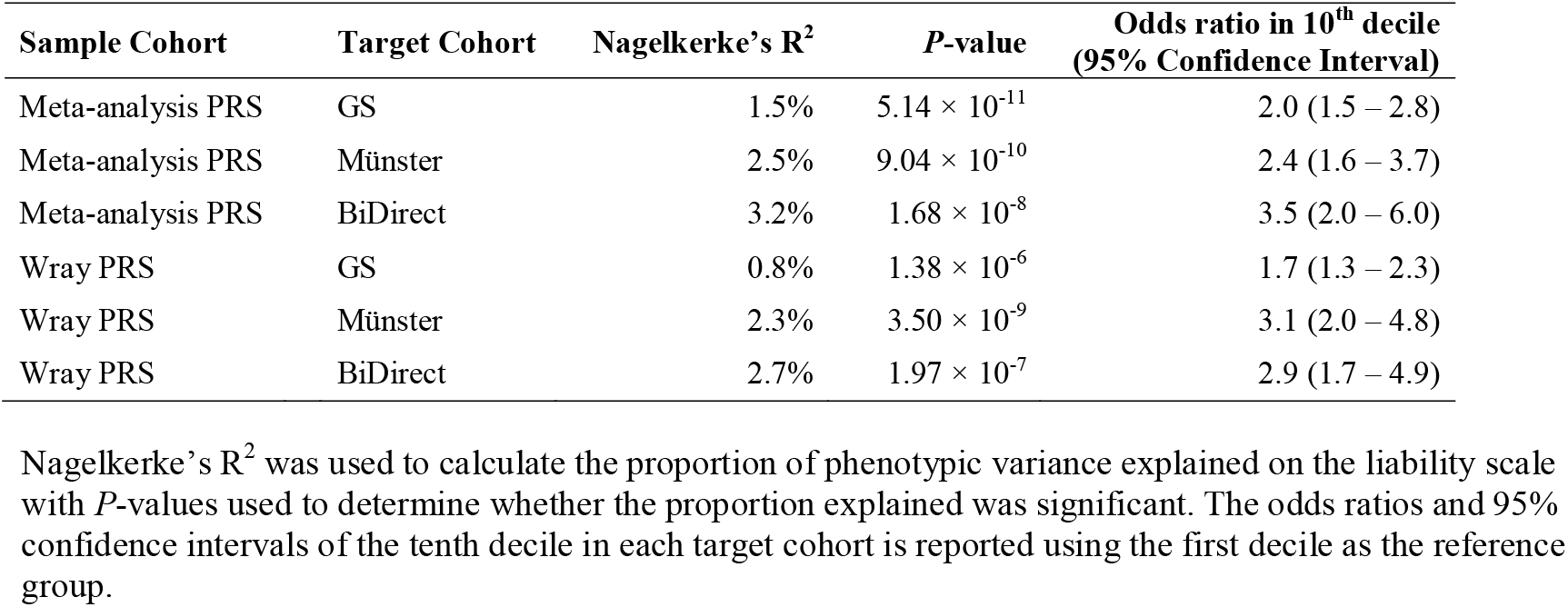
Out of sample prediction for depression using polygenic risk scores obtained from the current meta-analysis and from Wray, et al. ^9^ with Generation Scotland (GS), Münster and BiDirect as the target cohorts

| **Sample Cohort** | **Target Cohort** | **Nagelkerke's R<sup>2</sup>** | **P-value** | **Odds ratio in 10<sup>th</sup> decile**<br>**(95% Confidence Interval)** |
| --- | --- | --- | --- | --- |
| Meta-analysis PRS | GS | 1.5% | $5.14 \times 10^{-11}$ | 2.0 (1.5 – 2.8) |
| Meta-analysis PRS | Münster | 2.5% | $9.04 \times 10^{-10}$ | 2.4 (1.6 – 3.7) |
| Meta-analysis PRS | BiDirect | 3.2% | $1.68 \times 10^{-8}$ | 3.5 (2.0 – 6.0) |
| Wray PRS | GS | 0.8% | $1.38 \times 10^{-6}$ | 1.7 (1.3 – 2.3) |
| Wray PRS | Münster | 2.3% | $3.50 \times 10^{-9}$ | 3.1 (2.0 – 4.8) |
| Wray PRS | BiDirect | 2.7% | $1.97 \times 10^{-7}$ | 2.9 (1.7 – 4.9) |

Nagelkerke's R<sup>2</sup> was used to calculate the proportion of phenotypic variance explained on the liability scale with P-values used to determine whether the proportion explained was significant. The odds ratios and 95% confidence intervals of the tenth decile in each target cohort is reported using the first decile as the reference group.

A Manhattan plot of our meta-analysis results is provided in Figure 1 with a quantile-quantile plot provided in Supplementary Figure 1. Linkage Disequilibrium Score (LDSC) regression ^11^ produced a genomic inflation factor (λ_GC_) estimate of 1.63 with an intercept of 1.015 (0.011) prior to inflation correction, indicating that the inflation was due to polygenic signal and unlikely to be confounded by population structure. All of the 102 associated variants had the same direction of effect on depression across the three contributing studies and also within an independent replication sample of 1,306,354 individuals (Table 1; 414,055 cases and 892,299 controls). In the replication sample, 97 out of the 102 associated variants were nominally significant (*P* < 0.05) and 87 were significant after Bonferroni correction (α□=□0.05 / 102; *P* < 4.90 × 10^-4^). Further examination of the general directionality agreement of associated variants found in the contributing studies to the meta-analysis is provided in Supplementary Table 2 and the Supplementary Information. In summary, the direction of effect of depression variants in the previous studies was consistent with the current meta-analysis.

**Figure. 1.**
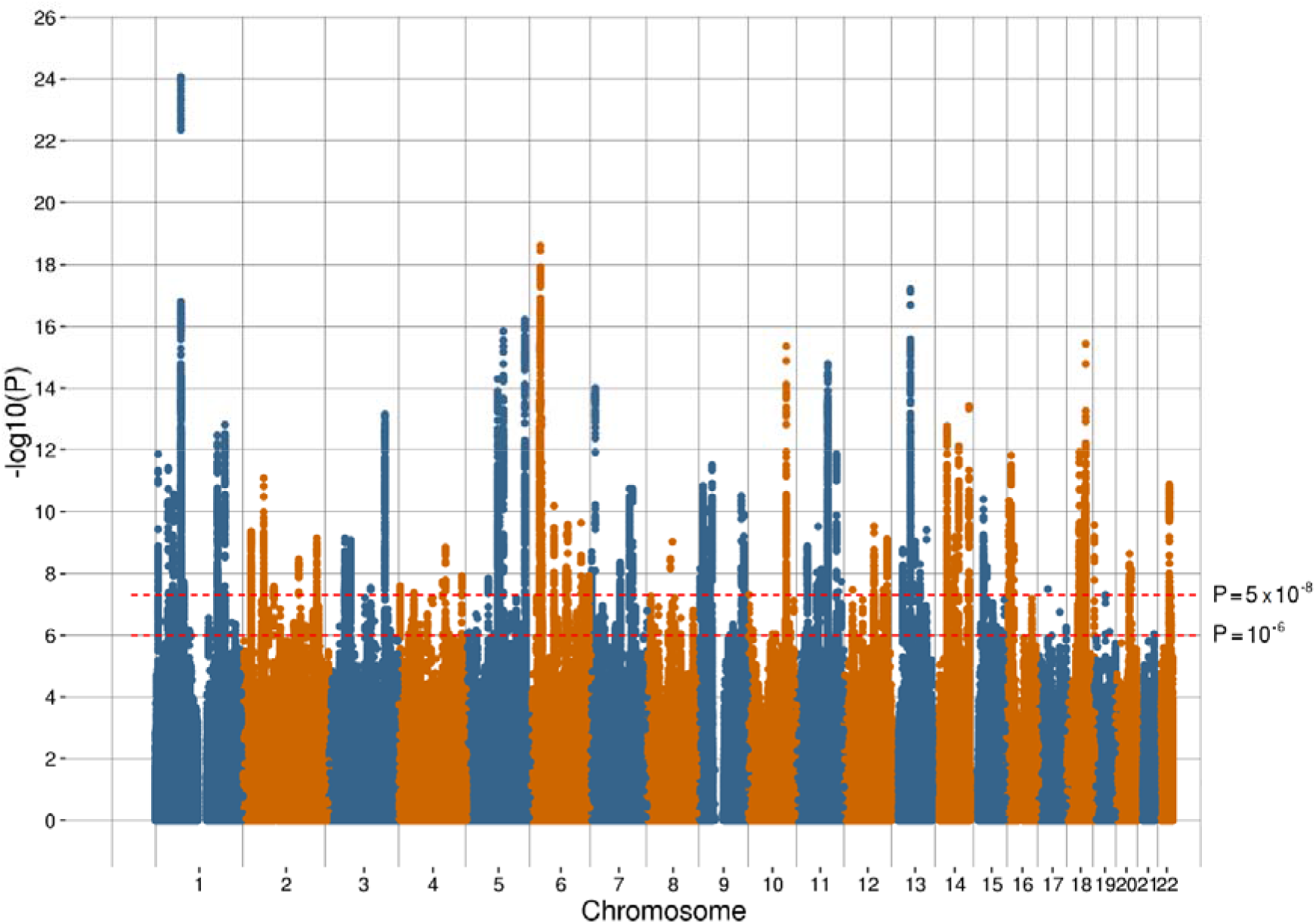
(A) Manhattan plot of the observed –log_10_ *P*-values of each variant for an association with depression in the meta-analysis (n = 807,553; 246,363 cases and 561,190 controls). Variants are positioned according to the GRCh37 assembly

Polygenic risk scores (PRS) were used to assess the predictive ability of the current genome-wide meta-analysis of depression within the clinically diagnosed MDD cohorts of Generation Scotland (GS; 975 cases and 5,971 controls), Münster (960 cases and 834 controls) and BiDirect (811 cases and 469 controls). PRS were also calculated using summary statistics from Wray, et al. ^9^ (Wray PRS) for comparison. Both the meta-analysis PRS and the Wray PRS were significantly associated with MDD in each of the three cohorts and the current meta-analysis PRS explained a greater proportion of the phenotypic variance in each target cohort compared to Wray PRS (Table 3 and Supplementary Figure 2).

**Table 3.** Gene pathways with a significant effect (*P*_corrected_ < 0.05) on depression identified through gene-set enrichment analysis in MAGMA ^22^

| Pathway | Number of genes | Beta | Standard error | P-value | $P_{\text{Corrected}}$ |
| --- | --- | --- | --- | --- | --- |
| GO_POSTSYNAPSE | 351 | 0.34 | 0.056 | $8.15 \times 10^{-10}$ | $4.82 \times 10^{-6}$ |
| GO_SYNAPSE | 707 | 0.219 | 0.040 | $2.47 \times 10^{-8}$ | $1.46 \times 10^{-4}$ |
| GO_NEURON_SPINE | 114 | 0.509 | 0.096 | $6.13 \times 10^{-8}$ | $3.63 \times 10^{-4}$ |
| GO_EXCITATORY_SYNAPSE | 181 | 0.405 | 0.077 | $8.29 \times 10^{-8}$ | $4.90 \times 10^{-4}$ |
| GO_SYNAPSE_PART | 571 | 0.225 | 0.044 | $1.24 \times 10^{-7}$ | $7.32 \times 10^{-4}$ |
| GO_BEHAVIOR | 489 | 0.245 | 0.049 | $2.32 \times 10^{-7}$ | $1.38 \times 10^{-3}$ |
| GO_COGNITION | 238 | 0.346 | 0.069 | $2.77 \times 10^{-7}$ | $1.64 \times 10^{-3}$ |
| GO_NEURON_PROJECTION | 883 | 0.18 | 0.036 | $3.39 \times 10^{-7}$ | $2.01 \times 10^{-3}$ |
| GO_MODULATION_OF_SYNAPTIC_TRANSMISSION | 283 | 0.298 | 0.061 | $4.25 \times 10^{-7}$ | $2.52 \times 10^{-3}$ |
| GO_REGULATION_OF_SYNAPSE_STRUCTURE_OR_ACTIVITY | 219 | 0.335 | 0.071 | $1.19 \times 10^{-6}$ | $7.02 \times 10^{-3}$ |
| GO_REGULATION_OF_SYNAPTIC_PLASTICITY | 133 | 0.412 | 0.089 | $1.95 \times 10^{-6}$ | 0.012 |
| GO_NEURON_PART | 1183 | 0.142 | 0.031 | $3.17 \times 10^{-6}$ | 0.019 |
| GO_DENDRITE | 420 | 0.229 | 0.052 | $4.76 \times 10^{-6}$ | 0.028 |
| GO_SINGLE_ORGANISM_BEHAVIOR | 367 | 0.244 | 0.056 | $5.86 \times 10^{-6}$ | 0.035 |
| Somatosensory Pyramidal Neurons | 1217 | 0.121 | 0.031 | $3.87 \times 10^{-5}$ | $2.71 \times 10^{-4}$ |

Pathways with a “GO” prefix were obtained from the Gene Ontology Consortium <sup>114</sup>. The Somatosensory Pyramidal Neurons gene-set was obtained from Skene, et al. <sup>23</sup>.

### Genetic correlations with depression

LDSC regression ^11^ was used to calculate pairwise genetic correlations (r_G_) to determine the extent of overlap in the genetic architectures across the three non-overlapping cohorts that contributed to our meta-analysis, i.e. between the UK Biobank, PGC_139k and 23andMe_307k depression analyses. There was a strong genetic correlation between each of these cohorts. The r_G_ between PGC_139k and 23andMe_307k was 0.85 (s.e = 0.03). The UK Biobank had a r_G_ of 0.87 (s.e = 0.04) with PGC_139k. A similar r_G_ was also found between UK Biobank and 23andMe_307k (0.85, s.e = 0.03). This was despite UK Biobank using a broader phenotype based on self-reported help-seeking behaviour compared to the self-declared clinical depression phenotype of 23andMe_307k and the primarily clinically obtained MDD phenotype of PGC_139k.

Depression is known to be comorbid with a wide range of other diseases and disorders^12^. To assess the shared genetic architecture between depression and many other traits, genetic correlations were calculated between our meta-analysed summary statistics of all three cohorts for depression and 234 behavioural and disease traits available via LD Hub ^13^ which implements LDSC regression. Of these behavioural and disease traits, 41 were significantly genetically correlated (*P*_FDR_ < 0.01) with our meta-analysis results after applying false discovery rate correction, see Supplementary Figure 3 and Supplementary Table 3. Significant genetic correlations with depression included schizophrenia (r_G_ = 0.32, s.e = 0.02), bipolar disorder (r_G_ = 0.33, s.e = 0.03), college completion (r_G_ = −0.19, s.e = 0.03), coronary artery disease (r_G_ = 0.13, s.e = 0.02), triglycerides (r_G_ = 0.14, s.e = 0.02), body fat (r_G_ = 0.16, s.e = 0.03), and waist-to-hip ratio (r_G_ = 0.12, s.e = 0.02). Many of these genetic correlations are similar to those reported by Wray, et al. ^9^ and Howard, et al. 5, including earlier age at menarche (r_G_ = −0.12, s.e = 0.02). However, a novel genetic correlation was observed for age at menopause (r_G_ = −0.11, s.e. 0.03), potentially indicating a shared genetic architecture between depression and earlier female reproductive life events. Additionally, novel genetic correlations were observed between depression and Crohn’s disease (r_G_ = 0.09, s.e. 0.03) and depression and an earlier age of smoking initiation (r_G_ = −0.21, s.e. 0.06).

### Causal relationships between depression and other traits

The genetic correlations between depression and other traits/disorders reported in the previous section may arise from genes with pleiotropic effects and biological influences across both traits. Alternatively, there may be a causal effect of depression on other traits (e.g. depression influencing triglyceride level) or from other traits causally influencing depression (e.g. triglyceride level leading to depression). To determine whether causal relationships exist between depression and the 41 genetically correlated traits in Supplementary Table 3, we used a bi-directional, two-sample Mendelian randomisation (MR) approach using MR-Base v0.4.9 ^14^ in R and an inverse-variance weighted (IVW) regression analysis. Where there was also evidence of variant heterogeneity additional sensitivity tests were conducted using an MR Egger test and a weighted median test (Supplementary Table 4).

A number of causal relationships were not tested because of sample overlap, which can lead to biased effect size estimates^15^, or where there were an insufficient number of instrumental genetic variables (nvariables < 30). A total of 33 causal effects were tested, of which 24 were for a causal effect of depression on another trait, and nine tests for a causal effect of another trait on depression. Directional horizontal pleiotropy, where a genetic variant has an effect on both traits but via differing biological pathways, can bias the estimates of causal effects between traits^16^. Using the MR Egger intercept test, there was no evidence (*P* ≥ 0.05) of directional horizontal pleiotropy for any of the 33 causal effects examined.

A putative causal effect of depression on neuroticism^17^ was detected using the IVW regression analysis MR test at the 1% significance threshold after false discovery rate (FDR) correction (beta = 0.146, s.e. = 0.039; *P*_FDR_ = 2.29 × 10^-3^; Supplementary Figure 4). However, there was also evidence of variant heterogeneity (*P* = 6.01 × 10^-3^), due to global horizontal pleiotropy, requiring additional sensitivity tests to examine the consistency of the effect. The additional sensitivity tests both had effects in the same direction as the IVW test (weighted median beta = 0.119, s.e. = 0.047; MR Egger beta = 0.050, s.e. = 0.235). There was also evidence of a putative causal effect of depression on ever vs. never smoked^18^ (beta = 0.285, s.e. = 0.077; *P*_FDR_ = 2.29 × 10^-3^; Supplementary Figure 5), with no evidence of variant heterogeneity (*P* = 0.14). Both the putative causal effects of depression on neuroticism and depression on ever vs. never smoked remained consistent (*P* < 6.46 × 10^-4^) in the ‘leave one variant out’ IVW analysis indicating that the observed effect was not driven by a single outlying variant.

Neuroticism was the only trait with a putative causal effect on depression using the IVW regression analysis MR test (beta = 0.366, s.e = 0.037; *P*_FDR_ = 2.63 × 10^-21^; Supplementary Figure 6). However, there was also evidence of variant heterogeneity (*P* = 9.62 × 10^-21^) requiring additional MR sensitivity testing. The weighted median test produced similar effect size and *P*-value to the IVW test (beta = 0.337, s.e. = 0.038; *P* = 1.94 × 10^-18^), but the MR Egger test had a large standard error and was in the opposite direction (beta = −0.128, s.e. = 0.271; *P* = 0.64). This putative causal effect remained consistent (*P* = 1.58 × 10^-21^) in the ‘leave one variant out’, IVW analysis indicating that the observed effect was not driven by a single variant.

The observed bi-directional relationship between depression and neuroticism could be confounded by non-independent instrumental variants across both tests, i.e. a region containing variants associated with depression was used to test for a causal effect on neuroticism and then variants in that same region were also used to test for a causal effect of neuroticism on depression. To account for any overlap, we identified and removed 15 instrumental variants from each bi-directional test where there was evidence of linkage disequilibrium (LD r^2^ > 0.1). The effect size for the IVW regression analysis MR test for depression on neuroticism was attenuated from 0.146 (s.e. = 0.039) to 0.112 (s.e. = 0.042) and the *P*-value was no longer significant after false discovery rate correction (*P*_FDR_ = 0.037). The effect size for the IVW regression analysis MR test for neuroticism on depression was attenuated from 0.366 (s.e. = 0.037) to 0.289 (s.e. = 0.042) and the *P*-value remained significant after false discovery rate correction (*P*_FDR_ = 1.34 × 10^-10^).

### Partitioning of the heritability component of depression

The estimate of the SNP-based heritability of depression within our meta-analysis was 0.089 (0.003) on the liability scale using LDSC regression ^11^. Heritability was then partitioned by calculating the proportion of heritability assigned to 24 functional categories divided by the proportion of variants in that category ^19^. This partitioning showed significant enrichment within conserved, intronic and H3K4me1 regions of the genome for the heritable component of depression (*P*_corrected_ < 0.05), see Supplementary Figure 7 and Supplementary Table 5. However, the estimates of enrichment for intronic (1.16×, s.e. = 0.05) and H3K4me1 (1.41×, s.e. = 0.13) regions were much smaller compared to the conserved regions (17.49×, s.e. = 1.68) of the genome.

Partitioning the heritability estimate by cell type enrichment (Figure 2A, and Supplementary Table 6) revealed that central nervous system (CNS) and skeletal muscle tissues were enriched (*P*_corrected_ < 0.05) for genetic variants contributing to the heritability of depression. Studies have reported altered histone modifications of skeletal muscle in response to exercise^20^ and suggested roles for skeletal muscle PGC-1α1^21^ in depression. The prominence of CNS enrichment led us to examine both brain regions (Figure 2B, Figure 2C, Supplementary Figure 8, and Supplementary Table 7) and brain cell types (Supplementary Figure 9, and Supplementary Table 8). There was significant enrichment (*P*_corrected_ < 0.05) of the anterior cingulate cortex, frontal cortex and cortex brain regions and neuron brain cells. The pseudo-coloring used in Figure 2C and Supplementary Figure 8 highlight in red the significance and effects sizes of the enriched regions of the brain associated with depression, respectively.

**Figure. 2.**
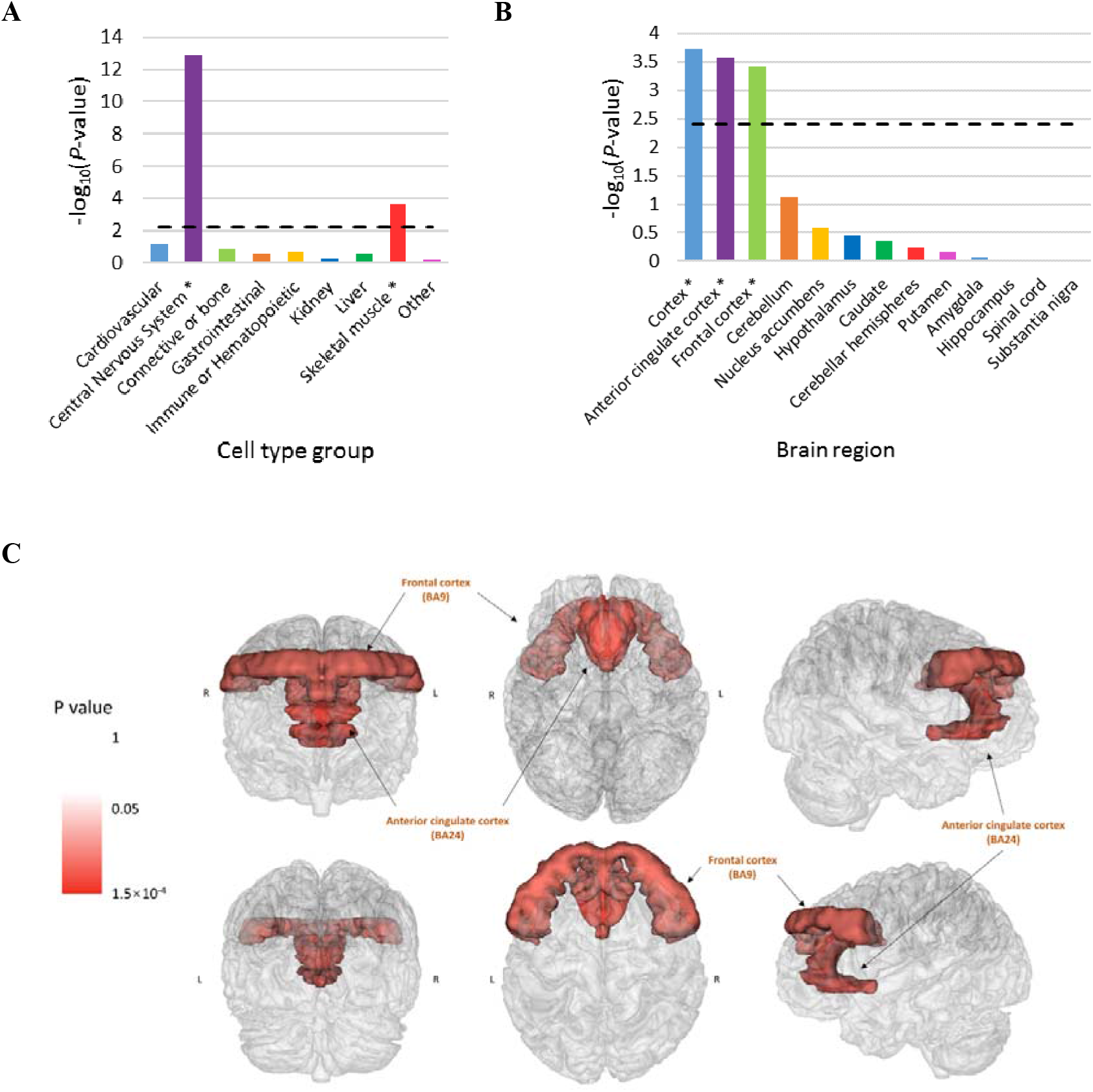
(A) Significance of cell type enrichment using a partitioned heritability approach. The dashed line represents statistical significance after Bonferroni correction (-log_10_*P* = 2.36) and * indicates significant enrichment for that cell type (B) Significance of enrichment estimates, based on genetic summary statistics, for brain regions using GTEx. The dashed line represents the Bonferroni cut-off for statistical significance (-log_10_*P* = 2.41) and * indicates significant enrichment for that brain region (C) Significant enrichment *P*-values, based on genetic summary statistics, for brain cell regions using GTEx overlaid on a physical representation of brain anatomy The pseudo-coloring used in Figure 2C highlights the *P*-values of the regions of the brain in red that were significantly enriched (*P* < 0.05) for depression variants

### Genes and gene-sets putatively associated with depression

The MAGMA ^22^ package was used to assess the aggregated genetic effects from the meta-analysis on 17,842 genes. This technique identified a total of 269 putative genes that were associated (*P* < 2.80 × 10^-6^) with depression, Supplementary Table 9. The most significant gene (*P* = 2.27 × 10^-19^) was Sortilin related VPS10 domain containing receptor 3 (*SORCS3*) on chromosome 10 (Supplementary Figure 10). The Neuronal Growth Regulator 1 (*NEGR1*) gene was associated with depression (*P* = 3.55 × 10^-15^) and there were two nearby independently-associated variants (rs2568958 and rs10890020) located in separate loci (Supplementary Figure 11 A-B). rs10890020 overlapped with the Long Intergenic Non-Protein Coding RNA 1360 (*LINC01306*) coding region which was not available for analysis in MAGMA. A further short (1.2 Mb) region along 18q.21.2 contained three independently-associated variants across two loci (rs62091461, rs12966052 and rs12967143; Supplementary Figure 12 A-C). These variants were close to the Transcription Factor 4 (*TCF4*) and the *RAB27B* genes which were both putatively associated with depression (*P* = 4.55 × 10^-16^ and *P* = 1.39 × 10^-9^, respectively). Further consideration of the genes putatively associated with depression is provided in the discussion.

To identify the biological pathways that are influenced by the putative genes associated with depression, gene-set analyses were performed using MAGMA ^22^. This method identifies the genes involved in each biological pathway and assesses whether there is evidence of enrichment for each pathway in depression using the *P*-values of each gene. We identified 14 significant putative gene-sets (*P*_corrected_ < 0.05) for depression (Table 3) using data from the Gene Ontology Consortium. Eight of these gene-sets were cellular components (areas where the genes were active) and were located in the nervous system: GO_POSTSYNAPSE, GO_SYNAPSE, GO_NEURON_SPINE, GO_EXCITATORY_SYNAPSE, GO_SYNAPSE_PART, GO_NEURON_PROJECTION, GO_NEURON_PART, and GO_DENDRITE. The cellular component gene-sets intimate a role in synapse function and excitatory mechanisms. The other six associated gene-sets were biological processes: GO_BEHAVIOR, CO_COGNITION, GO_MODULATION_OF_SYNAPTIC_TRANSMISSION, GO_REGULATION_OF_SYNAPSE_STRUCTURE_OR_ACTIVITY, GO_MODULATION_OF_SYNAPTIC_PLASTICITY, and GO_SINGLE_ORGANISM_BEHAVIOR. The gene overlap between these gene-sets (Supplementary Table 10) suggests that the gene-sets generally fall within two clusters. One gene-set cluster typically relates to synaptic structure and activity while the other gene-set cluster relates to the response or behaviour to stimuli. We also identified a significant gene-set (*P* = 3.87 × 10^-5^) containing somatosensory pyramidal neurons using brain cell-type data from Skene, et al. ^23^ (Table 3).

### Drug - gene interactions

The 269 putative genes that were identified as being significantly associated (*P* < 2.80 × 10^-6^) with depression using MAGMA ^22^ were examined for known interactions (including agonistic, partial agonistic, antagonistic, modulating, binding and blocking interactions) with prescribed medications in the Drug Gene Interaction Database (dgidb.genome.wustl.edu) v3.0 ^24^. A total of 560 interactions were identified between 57 genes and 514 drugs (see Supplementary Table 11). Anatomical Therapeutic Chemical (ATC) classifications were available for 220 of these drugs which belonged to 54 different second level ATC classes and which interacted with 37 genes, Figure 3. The greatest number of drug - gene interactions (ninteractions = 47) were observed between psycholeptics (N05, which includes antipsychotics and anxiolytics) and dopamine receptor D2 (*DRD2*).

**Figure. 3.**
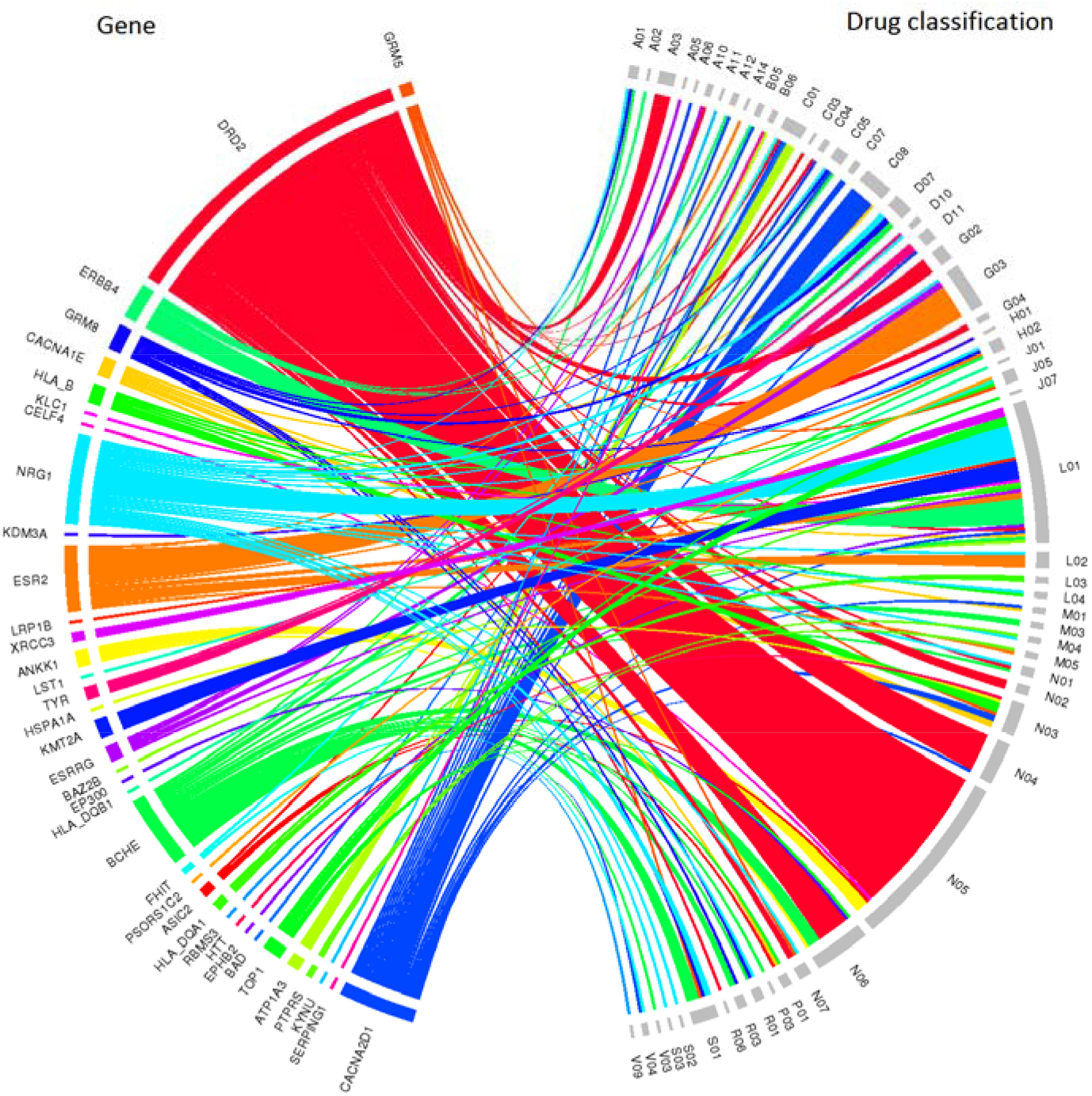
Chord diagram of genes significantly associated (*P* < 2.80 × 10-6) with depression and the second level Anatomical Therapeutic Chemical classifications to which interacting drugs have been assigned. The width of each line is determined by the number of drugs known to interact with each gene. The genes are ordered by significance with depression with those most significantly associated located at the top of the diagram

## Discussion

In this study we report a meta-analysis of 807,553 individuals (246,363 cases and 561,190 controls) using summary statistics from three independent studies of depression, Hyde, et al. ^8^ (23andMe_307k), Howard, et al. ^5^ (UK Biobank) and Wray, et al. 9 (PGC_139k). The meta-analysis examined 8,098,588 variants and identified 102 independently segregating variants associated (*P* < 5 × 10^-8^) with depression. These 102 variants were assessed in an independent replication sample of 1,306,354 individuals (414,055 cases and 892,299 controls) and 87 variants were significant in that sample after multiple testing correction. The estimate of the genome-wide SNP-based heritability on the liability scale was 0.089 (s.e. = 0.003), indicating that the analysed depression phenotype had a significant genetic component. All of the 102 variants associated with depression had an equivalent direction of allelic effect across the three studies^5, 8, 9^ that contributed to the meta-analysis and the replication sample (Supplementary Table 1), suggesting that these variants represent robust associations with depression. Our gene-based analyses revealed pathways relating to neurotransmission and response to stimuli, with the central nervous system also found to be enriched when performing heritability partitioning. The partitioning of the heritability for depression also highlighted the importance of the cortical regions of the brain and further research focussed on the regions and biological pathways detected is warranted. We detected 269 genes putatively associated with depression and demonstrated the utility of investigating their interaction with pharmacological treatments.

The three studies^5, 8, 9^ contributing to the meta-analysis were based on different depression phenotypes; nevertheless, there were strong genetic correlations (> 0.85) between them. PGC_139k used a predominately clinically ascertained diagnosis of major depression and the observed genetic correlations indicate that each study was likely to be capturing a similar underlying genetic architecture, despite the use of different diagnostic instruments. Therefore, larger population-based samples, where the timescales and costs of obtaining a clinically diagnosis would be prohibitive, can contribute to our understanding of the genetic architecture of the disease. The meta-analysis also provides the opportunity to assess the general directionality agreement of variants that have previously been associated with depression (Supplementary Information and Supplementary Table 2). The examination of previous findings between studies and the current meta-analysis suggests that there is good degree of directionality agreement of causal effects of depression when using studies with over 100,000 participants. However, there is likely to be a need to continue to ascertain clinically ascertained MDD cohorts to ensure validity of the larger cohorts with broader phenotyping.

There is an ever growing body of evidence that there are shared genetic components between behavioural and psychiatric disorders^25–27^. Using the meta-analysed results in the current study this was evidenced by significant genetic correlations for depression with neuroticism, anorexia nervosa, attention deficit hyperactivity disorder (ADHD), schizophrenia, and bipolar disorder. The MR analysis also identified a putative bi-directional casual effect between depression and neuroticism. Removing regions which were used to test the effect in both directions indicated that a putative effect remained only for neuroticism on depression. This uni-directional effect does make more sense intuitively as neuroticism is a stable trait whereas depression is a more episodic condition.

Examining significant genes that overlap between the current meta-analysis of depression and the neuroticism studies of both Luciano, et al. ^28^ and Okbay, et al. ^29^ revealed putative associations with *DRD2*, CUGBP Elav-like family member 4 (*CELF4*) and ELAV like RNA binding protein 2 (*ELAVL2*). Dopamine acts as a neurotransmitter in the brain with *DRD2* encoding the dopamine receptor D2 subtype. Genetic variation around the *DRD2* gene has been linked to differences in structural connectivity between the basal ganglia structure with the frontal cortices^30^. Supporting evidence of the importance of the cortical brain regions was provided by the heritability partitioning of the brain regions (Figures 2B and 2C) with significant enrichment of the anterior cingulate cortex, frontal cortex and cortex brain regions. *DRD2* is also associated with mood modulation and emotion processing^31^ and is also commonly reported in association studies of schizophrenia^32, 33^. *DRD2* was included in all 14 of the putative Gene Ontology Consortium gene-sets identified (but not the somatosensory pyramidal neurons gene-set) and has been identified in previous studies of depression^5, 9, 34^. *CELF4* plays a key role in coordinating the synaptic function in excitatory neurons^35^ with dynamic changes in expression during brain development^36^ and deletions in the surrounding region (18q.12.2) associated with autism spectrum disorder^37^ and developmental and behavioural disorders^38^. *ELAVL2* potentially aids in the regulation of gene expression pathways in human neurodevelopment ^39^ and disruption of related pathways may be a factor in neurodevelopmental disorders^40^. Improving our understanding of the genetic similarities and differences between neuroticism and depression may provide a route to determining the biological aspects that underpin a more permanent personality trait or a depressive state. These genetic similarities and differences also provide opportunities for phenotypic stratification and warrants further investigation in future studies.

The current study further reaffirms the genetic correlation between schizophrenia and depression observed by Wray, et al. ^9^. The current gene-based analysis identified the vaccinia-related kinase 2 (*VRK2*) gene as putatively associated with depression. Extensive research has been conducted examining the link between *VRK2* and schizophrenia, with the association replicated in both European ^41, 42^ and Asian ^43^ populations. Additionally, associations have been found between schizophrenia and the arginine and serine rich coiled-coil 1 (*RSRC1*) ^44^ and myocyte enhancer factor 2C (*MEF2C*) ^32, 45^ genes, both of which we found to be putatively associated with depression. Hyde et al. ^8^ reported that *RSRC1* and *MEF2C* were also close to the genetic variants that they detected, and they highlighted *MEF2C*’s role in regulation of synaptic function and central nervous system phenotypes. *MEF2C* was also included in ten of the 15 putative gene-sets identified for depression. The *TCF4* gene coding region (Supplementary Figure 12) is also noteworthy as it contained two independently-associated variants for depression and has been identified in previous studies of depression ^5, 9, 46^. *TCF4* is involved in the synaptic plasticity ^47^ and the excitability of prefrontal neurons ^48^ and has been implicated in other psychiatric disorders^49^, including schizophrenia ^32^.

ADHD is a neurodevelopmental disorder, typically diagnosed during early childhood (age 4 to 6) and is associated with an increased risk of depression during adolescence ^50^. The current study demonstrated that there was a significant genetic correlation between ADHD and depression. The cadherin 13 (*CDH13*) gene, which codes for cell adhesion molecules, was found to be putatively associated with depression and has also been implicated in ADHD ^51, 52^ and specifically hyperactive and impulsive symptoms ^53^. *CDH13* was included in the putative GO_NEURON_PART and GO_NEURON_PROJECTION gene-sets. A further gene involved in cell adhesion, astrotactin 2 (*ASTN2*), which we found to be putatively associated with depression is also involved in ADHD aetiology ^52, 54^ and plays a role in neuronal development in the brain ^55^. Dopamine transmission may also underlie ADHD and depression with both the *DRD2* and ankyrin repeat and kinase domain containing 1 (*ANKK1*) genes implicated in our analysis of depression and in studies of ADHD ^56, 57^.

Age of menarche has been found to be phenotypically and genetically correlated with depression ^9, 58, 59^, with a causal effect of earlier age at menarche on depressive symptoms also reported ^60^. The current meta-analysis identified a significant genetic correlation between age of menarche and depression, and nominal evidence of a bi-directional causal effect. The lin-28 homolog b (*LIN28B*) gene was significant in our meta-analysis and has frequently been associated with age of menarche ^61–63^. The biological mechanisms that underlie the association of *LIN28B* with both age of menarche and depression remain unclear; however, *LIN28B* has been shown to be involved in regulating cell pluripotency ^64^ and developmental timing ^65^, and through its mediation of Lethal-7 miRNA has been implicated in inflammation and immune response ^66^. The current analysis also identified a novel genetic correlation between age of menopause and depression which may share a similar biological mechanism to that of age of menarche.

A number of studies, using different methodologies, have examined causal relationships between depression and smoking with smoking increasing the risk of depression ^67, 68^, a bi-directional effect ^69^ and no effect ^70–73^ reported. In the current analysis there was an insufficient number of instrumental variants to test the effect of smoking on depression, but there was evidence of a genetic correlation between depression and cigarette smoking as well as a causal effect of depression on smoking. Smoking has been reported to have an anxiolytic and antidepressant effect ^74, 75^ and this may explain why we observe a causal effect of depression on smoking; however, as reported by Munafò, Hitsman, Rende, Metcalfe and Niaura ^76^ the relationship is likely to be complex and requires further investigation.

### Drug - gene interactions

Examining the number of interactions between the genes associated with depression and the second level ATC drug classifications demonstrates that there are currently available pharmaceutical treatments that may target the genetic component of depression. Most notable were the large number of interactions between the *DRD2* gene and the N05 drug classification, which primarily comprises typical and atypical antipsychotics. The dopaminergic system has been previously implicated in depression, particularly the symptoms of anhedonia and decreased motivation, and antidepressant action has been reported for dopamine reuptake inhibitors and D2-receptor agonists in animal models of depression ^77^. Furthermore, the dopamine and noradrenaline reuptake inhibitor bupropion is licensed for the treatment of depression ^78^.

We also identified genes with associated medications which, although not aimed at treating depression, may provide unpredicted drug benefits or adverse effects for those with the disorder. The Neuregulin 1 (*NRG1*) receptor ErbB4, found on GABAergic neurones, has been identified as a potentially druggable target for depression, anxiety and schizophrenia ^79^. There is interest in developing agents, such as basimglurant and fenobam, as antidepressants and anxiolytics that target glutamatergic receptors ^80^. Estrogen Receptor 2 (*ESR2*) has previously been implicated in antidepressant action through up-regulation by 3β-hydroxysteroid dehydrogenase ^81^. Estrogenic compounds have also been found to have antidepressant effects in rodent models of depression ^82^.

Further, our work highlights other potential druggable genes associated with depression which are not, to our knowledge, currently associated with antidepressant treatment or mood-associated adverse effects. These include the R-type calcium channel gene, Calcium Voltage-Gated Channel Subunit Alpha1 E (*CACNA1E*), and the nucleosome associated gene, Lysine Methyltransferase 2A (*KMT2A*) ^83^.

An intriguing omission among the depression-associated genes identified in our study are genes linked with the serotonergic system, such as the serotonin transporter *SLC6A4*; the G protein subunit *GNB3*; the serotonin receptor *HTR2A* and tryptophan hydroxylase (*TPH2*). This is surprising, as interaction with the serotonergic system forms the basis of most antidepressant treatments. Our finding could indicate a functional separation between genetic pathways of depressive disease and pathways of antidepressant treatment. Thus, serotonin-associated genes, while potentially relevant for predicting efficacy and adverse effects of serotonergic antidepressants, may not be directly associated with the aetiology of depression itself (or at least that which is determined by common genetic variation identified in GWAS). Indeed, a recent review of the genetics studies of depression has remarked upon the failure to demonstrate association with depression for serotonergic and other popular candidate genes ^84^. It may also be that the pathways of depression and serotonergic antidepressant effect are separate but entwine through common intermediary genes. One such candidate is *NRG1*, identified here in depression, and also in a recent meta-analysis of antidepressant response ^85^. These findings suggest that there is potentially a need to concurrently model a range of ‘omics data, including genomics, epigenomics, and transcriptomics to gain further understanding of depression pharmacology.

The principal strength of this study is the increased power obtained from the analysis of three large independent cohorts. This has allowed the examination of the effects of variants and regions that have been identified previously to determine whether they maintain an effect on depression. We have used variant-based analyses to calculate heritability enrichment across different brain regions and then also applied a complimentary approach using the genes assigned to functional gene-sets, with both methods highlighting the importance of the prefrontal brain regions. Our study highlights a number of potential gene targets for drug repositioning; however, due to the causal heterogeneity of depression ^86^, it is likely that stratifying depression will lead to clearer distinction between depression subtypes and potential treatments.

## Conclusion

This study describes a meta-analysis of the three largest depression cohorts (total n = 807,553, with 246,363 cases and 561,190 controls) currently available. The meta-analysis identified 102 independently-segregating genetic variants associated with depression in 101 loci and demonstrated a consistency of effect directionality in a large replication sample (total n = 1,306,354, with 414,055 cases and 892,299 controls) and across the three contributing studies allowing greater confidence in the findings. The heritability enrichments and gene-set analysis both provided evidence for the role of prefrontal brain regions in depression and the genes identified contribute to our understanding of biological mechanisms and potential drug targeting opportunities. These findings advance our understanding of the underlying genetic architecture of depression and provide novel avenues for future research.

## Online Methods

To conduct our analyses we included data from three previous studies of depression ^5, 8, 9^. For the UK Biobank study ^5^ slightly different quality control was applied and the data were reanalysed. For the other two studies the summary statistics were obtained directly from the respective authors. Further information regarding each of these cohorts is provided below.

### UK Biobank

The population-based UK Biobank cohort ^87^ consists of 501,726 individuals with genome-wide data for 93,095,623 autosomal genetic variants imputed using the HRC and UK10K reference panels ^88^. We used the variants from the HRC reference panel to conduct a 4-means clustering approach and used the first two principal components to identify a genetically homogenous subgroup of 462,065 individuals. We next removed 131,790 individuals that had a shared relatedness up to the third degree that were identified by UK Biobank based on kinship coefficients (> 0.044) calculated using the KING toolset ^89^. For these 131,790 removed individuals we then calculated a genomic relationship matrix and identified one individual to be reinstated from within each related group that had a genetic relatedness less than 0.025 with all other participants which allowed us to add an additional 55,745 individuals back into our sample. We then used a checksum based approach ^90^ to identify and exclude 954 individuals from within the UK Biobank cohort that overlapped with the Major Depressive Disorder Working Group of the Psychiatric Genomics Consortium (PGC) cohorts analysed by Wray, et al. ^9^. This was possible for a total of 30 out of 33 cohorts that make up the PGC analysis due to the availability of genetic data. We also removed those UK Biobank individuals with a variant call rate < 90% or that were outliers based on heterozygosity, or variants with a call rate < 98%, a minor allele frequency < 0.005, those that deviated from Hardy-Weinberg equilibrium (*P* < 10^-6^), or had an imputation accuracy score < 0.1, leaving a total of 10,612,809 variants for 371,437 individuals.

Within UK Biobank we used the broad definition of depression ^5^ with more detailed phenotypic information available in that paper. In summary, case and control status of broad depression was defined by the participants’ response to the questions ‘Have you ever seen a general practitioner for nerves, anxiety, tension or depression?’ or ‘Have you ever seen a psychiatrist for nerves, anxiety, tension or depression?’. Exclusions were applied to participants who were identified with bipolar disorder, schizophrenia, or personality disorder using self-declared data following the approach of Smith, et al. ^91^ as well as prescriptions for antipsychotic medications. This provided a total of 127,552 cases and 233,763 controls (n = 361,315, prevalence = 0.353) for analysis.

### Statistical Analysis of UK Biobank

To conduct the association analysis within UK Biobank we used BGENIE v1.1 ^87^ to assess the effect of each genetic variant using a linear association test:

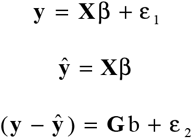

where **y** was the vector of binary observations for each phenotype (controls coded as 0 and cases coded as 1). β was the matrix of fixed effects, including sex, age, genotyping array, and the first 8 principal components and **X** was the corresponding incidence matrices. (**y − ŷ**) was a vector of phenotypes residualized on the fixed effect predictors, **G** was a vector of expected genotype counts of the effect allele (dosages), b was the effect of the genotype on residualized phenotypes, and ε_1_ and ε_2_ were vectors of normally distributed errors. The effect sizes and standard errors were transformed to the logistic scale by dividing each value by *ρ*(1- *ρ*), where *ρ* is the trait prevalence (0.353).

### 23andMe

We obtained the genome-wide association study results from the discovery 23andMe subset (23andMe_307k) from the Hyde, et al. ^8^ analysis. Phenotypic status was based on responses to web-based surveys with individuals that self-reported as having received a clinical diagnosis or treatment for depression classified as cases. This provided a total of 75,607 cases and 231,747 controls (n = 307,354, prevalence = 0.25). We excluded variants with an imputation accuracy threshold < 0.6 and with a minor allele frequency < 0.005, which left a total of 8,995,180 variants.

### Major Depressive Disorder Working Group of the Psychiatric Genomics Consortium (PGC)

Wray et al. conducted the largest meta-analysis of MDD to date ^9^, utilising European-ancestry PGC cohorts with an emphasis placed on obtaining clinically-derived phenotypes for MDD. Their meta-analysis included the 23andMe_307k discovery cohort ^8^ and a previous release of the UK Biobank data ^92^. We therefore obtained the summary statistics from their meta-analysis of major depression with the 23andMe_307k and the previous UK Biobank cohorts removed (PGC_139k). This provided a total of 12,149,399 variant calls for 43,204 cases and 95,680 controls (n = 138,884, prevalence = 0.31). We excluded variants with an imputation accuracy threshold < 0.6 or a minor allele frequency < 0.005, which left a total of 10,365,555 variants.

### Meta-analysis of genome-wide association study summary statistics

We used Metal ^93^ to conduct an inverse variance-weighted meta-analysis of the summary statistics from the three studies (using the log of the odds ratios and the standard errors of the log of the odds ratio), conditional on each variant being available in all three of the contributing cohorts. This provided 8,098,588 genetic variants and up to 246,363 cases and 561,190 controls (n = 807,553) for the meta-analysis. Linkage disequilibrium score (LDSC) regression intercepts ^11^ were used for genomic inflation control of the three contributing cohorts and the final meta-analysis results. Clumping and merging were used to identify the basepair positions of loci containing depression-associated variants. Clumping of the meta-analysis results was conducted using Plink v1.90b4 ^94^ applying the following parameters: –clump-p1 1e-4 –clump-p2 1e-4 –clump-r2 0.1 –clump-kb 3000, with merging of the clumped loci conducted using bedtools ^95^. A conditional analysis ^10^ was used to identify independently-segregating variants in each of the merged loci that was genome-wide significant, using the linkage disequilibrium structure of UK Biobank as a reference panel. Genome-wide statistical significance was defined as *P* < 5 × 10^-8^, with the meta-analysis results from Metal reported. Due to the complexity of the major histocompatibility complex region only the most significant variant in that region is reported.

### 23andMe replication sample

The variants that were significant (*P* < 5 × 10^-8^) in the meta-analysis, described above, were replicated in an independent sample of 1,306,354 unrelated individuals (414,055 cases and 892,299 controls) from 23andMe, Inc. Individuals within this replication sample were unrelated to individuals within 23andMe_307k. Detailed information regarding this replication sample is provided in the Supplementary Information. In summary, imputed genetic data was obtained for an unrelated European sample and the variants identified as genome-wide significant (*P* < 5 × 10^-8^) in the meta-analysis were analysed using logistic regression. The phenotype used in the 23andMe replication sample was the same used within the 23andMe_307k cohort, described above, with the responses to web-based surveys used to classify individuals that self-reported as having received a clinical diagnosis or treatment for depression classified as cases. We used Metal ^93^ to conduct an inverse variance-weighted meta-analysis of the significant variants (*P* < 5 × 10^-8^) identified in the previous paragraph (using UK Biobank, 23andMe_307 and PGC_139k) and the 23andMe replication cohort.

### Polygenic risk score analysis

Polygenic risk scores (PRS) were created using Plink v1.90b4 ^94^ and evaluated in the Generation Scotland: the Scottish Family Health Study (GS) ^96^, Münster ^9^ and BiDirect cohorts ^97^ using the same method described in Wray, et al. ^9^. Two polygenic risk scores were created: one using weightings from the current meta-analysis and one using weightings from a previous association study of major depression conducted by Wray, et al. ^9^ (Wray PRS). To create independent SNP clumping was applied using a linkage disequilibrium r^2^ of < 0.1 and a 500kb sliding window. PRS were calculated using *P*-value thresholds of ≤ 5 × 10^-8^, ≤ 1 × 10^-6^, ≤ 1 × 10^-4^, ≤ 0.001, ≤0.01, ≤0.05, ≤0.1, ≤ 0.2, ≤ 0.5 and the full model including all SNPs (*P* ≤ 1). Nagelkerke’s R^2^ was used to calculate the proportion of phenotypic variance explained on the liability scale. PRS were split into deciles and the odds ratio for MDD in each decile calculated using the 1st decile as the reference group.

GS is a family and population-based cohort of 20,195 participants genotyped on the Illumina OmniExpress BeadChip (706,786 SNPs). The raw genotype data in GS underwent quality control so that individuals with a call rate < 98%, SNPs with a genotyping rate < 98%, a minor allele frequency < 1% and a Hardy-Weinberg equilibrium *P*-value < 10^-6^ were removed from the dataset and then imputed ^98^, leaving a total of 19,997 GS individuals and 8,633,288 SNPs. Participants underwent a clinical diagnosis of MDD using the Structured Clinical Interview for the Diagnostic and Statistical Manual of Mental Disorders ^99^ with further information on the phenotype used provided in Fernandez-Pujals, et al. ^100^. An unrelated sample was selected by removing one individual from each pair that shared a genomic relationship > 0.025, leaving a total of 975 cases and 5,971 controls. To calculate the PRS, GS was removed from the current meta-analysis and the Wray PRS prior to calculated single nucleotide polymorphism (SNP) weights which were filtered so that they only contained SNPs that overlapped between the current meta-analysis PRS and the Wray PRS. The first five principal components in GS were fitted to account for population stratification. The best thresholds for the meta-analysis PRS (*P* ≤ 0.05; 62,519 SNPs) and Wray PRS (*P* ≤ 0.001; 5,941 SNPs) were each used to test for an association with MDD in GS.

The Münster cohort is described in Wray, et al. ^9^ and although this cohort was not part of their meta-analysis it was used for out of sample prediction using polygenic risk scores. In summary, the Münster cohort is a clinically ascertained sample with 960 MDD inpatient cases and 834 screened controls. The quality control procedures and the genome-wide association analysis of this cohort was conducting the same pipeline as used in Wray, et al. ^9^. The best thresholds for the meta-analysis PRS (*P* ≤ 0.01; 21,115 SNPs) and Wray PRS (*P* ≤ 0.05; 62,166 SNPs) were each used to test for an association with MDD in the Münster cohort.

The BiDirect cohort is a prospective observational study established to investigate the relationship between depression and cardiovascular disease ^97^. Cases were recruited from psychiatric and psychosomatic hospitals and residential psychiatrists’ practices in and around Münster and were required to be between the ages of 35 and 65 and had to be receiving in-patient or out-patient treatment for acute depression. A detailed description of the diagnostic criteria used in this cohort is provided by Teismann, et al. ^97^. Controls were randomly ascertained from the local population. The quality control procedures and genome-wide association analysis of this cohort was also conducting the same pipeline as used in Wray, et al. ^9^. There were a total of 811 acute depression cases and 469 controls used to calculate the proportion of variance explained using polygenic risk scores in the BiDirect cohort. The best thresholds for the meta-analysis PRS (*P* ≤ 0.5; 24,964 SNPs) and Wray PRS (*P* ≤ 0.05; 62,144 SNPs) were each used to test for an association with MDD in the BiDirect cohort.

### Genetic Correlations

We calculated genetic correlations using LDSC regression ^11^ and the online software LD hub (http://ldsc.broadinstitute.org/) ^13^. LDSC regression leverages linkage disequilibrium (LD) information for each genetic variant such that the χ^2^ statistic for that variant includes the effects of all loci in LD with that variant. This can be extended to the analysis of genetic correlations between traits if the χ^2^ statistic is replaced with the product of 2 z-scores from 2 traits of interest ^101^. Genetic correlations were calculated between the three datasets that contributed to the meta-analysis. Genetic correlations were also calculated between our meta-analysed results (23andMe_307k, UK Biobank and PGC_139k) for depression and 234 behavioural and disease related traits. *P*-values were false discovery rate (FDR) adjusted^102^ and correlations reported if *P*_FDR_ < 0.01.

### Mendelian randomization

We used Mendelian randomization (MR) to assess whether causal effects exist between depression and a number of other traits and disorders. MR uses genetic variants as a proxy for environmental exposures, assuming that: i) the genetic variants are associated with the exposure; ii) the genetic variants are independent of confounders in the exposure-outcome association; iii) the genetic variants are associated with the outcome only via their effect on the exposure, i.e. there is no horizontal pleiotropy whereby the variants affect both exposure and outcome independently. Individual genetic variants may be weak instruments for assessing causality, particularly if they have only small effect sizes. Using multiple genetic variants can increase the strength of the instrument, but also increases the risk of violating the MR assumptions.

We performed bidirectional, two-sample MR between our meta-analysis results for depression and all available traits which had a significant genetic correlation with depression (identified in the previous section). Traits directly related to or including depression (major depressive disorder, depressive symptoms and PGC cross disorder) were excluded due to potential bias. The genetic instruments for depression consisted of the independent, genome-wide significant variants, their effect sizes and standard errors, as estimated in our genome-wide meta-analysis. Summary statistics from genome-wide association studies for the other traits were sourced from either publicly available datasets or from the MR-Base database ^14^. Overlapping datasets for the exposure and the outcome can lead to bias and inflation of causal estimates ^15^. To mitigate this, when the source of the other trait included UK Biobank, 23andMe or any of the studies that contributed to PGC_139k, these studies were removed from our meta-analysis of depression; for example, the neuroticism trait from van den Berg, et al. ^17^ was assessed against a meta-analysis of depression using UK Biobank and 23andMe_307k only. Where UK Biobank, 23andMe and PGC_139k were all included in the genome-wide association study of the other trait then an alternative study was sought for that other trait.

All analyses were performed using the MR-Base “TwoSampleMR” v0.4.9 package^14^ in R. To avoid bias in the MR estimates due to linkage disequilibrium (r^2^), clumping was applied using the “clump_data” function with an r^2^ < 0.001. Genetic variants were required to be available in both the exposure and outcome traits and were harmonised using the default parameters within the TwoSampleMR package. Following this harmonisation, we only examined causal relationships where there were at least 30 instrumental genetic variables.

Directional horizontal pleiotropy, where a genetic instrument has an effect on an outcome independent of its influence on the exposure, can be a problem in MR analysis, particularly when multiple genetic variants of unknown function are used. We therefore firstly tested for directional horizontal pleiotropy using the MR Egger intercept test, as previously described by Hagenaars, Gale, Deary and Harris ^103^. If the MR Egger intercept test had a significant *P*-value (*P* < 0.05) then it was excluded from the analysis. However, no tests were excluded due to directional horizontal pleiotropy.

The second analysis conducted was a variant heterogeneity test for global horizontal pleiotropy. Variant heterogeneity is an important metric, but high heterogeneity doesn’t necessarily mean bias or unreliable results; for example, every instrumental variable could have horizontal pleiotropic effects but if they have a mean effect of 0 then there will be no bias, just larger standard errors due to more noise. For analyses that had evidence of high variant heterogeneity (*P* < 0.05), additional sensitivity MR tests were conducted. The sensitivity tests that were used were the MR Egger test and the weighted median test to examine whether the effect estimate was consistent.

The principal MR test of a causal effect was conducted using inverse-variance weighted (IVW) regression. This method is based on a regression of the exposure and the outcome which assumes the intercept is constrained to zero, and produces a causal estimate of the exposure-outcome association. Where there was no evidence of global horizontal pleiotropy (*P* ≥ 0.05) from the second analysis (see previous paragraph), a FDR adjusted^102^ *P*-value < 0.01 from the IVW test was required for evidence of a causal effect. Where there was evidence of global horizontal pleiotropy (P < 0.05) from the second analysis, additional evidence was also sought from the sensitivity tests (MR Egger test and the weighted median test). To ensure that a causal effect was not driven by a single variant a ‘leave one variant out’ IVW regression analysis was conducted with the least significant observed *P*-value used to assess whether significance was maintained. We tested the causal effect of depression on 24 other traits, and the causal effect of 9 other traits on depression.

### Partitioned heritability analyses

We used stratified LDSC regression to estimate the SNP-based heritability of depression, using the sample prevalence as the population prevalence (0.302), and then examined the heritability of partitioned functional categories ^19^. This method assigns variants into 24 functional categories and then evaluates the contribution of each category to the overall heritability of a trait. A category is enriched for heritability if the variants with high LD to that category have elevated χ^2^ statistics. The 24 categories are described in full in the Finucane et al. paper ^19^. Briefly, they include genetic annotations from ReqSeq gene models ^104^, transcription factor binding sites from ENCODE ^104, 105^, Roadmap epigenomics annotations ^106^, super-enhancers ^107^, evolutionarily conserved regions ^108^ and FANTOM5 enhancers ^109^. Heritability enrichment is defined as the proportion of heritability assigned to a functional category divided by the proportion of variants in that category. Cell-type specific annotations were also analysed for variants in four histone marks: H3K4me1, H3K4me3, H3K9ac and H3K27ac and these were grouped into 9 cell type groups: central nervous system (CNS), cardiovascular, connective tissue or bone, gastrointestinal, immune or hematopoietic, kidney, liver, skeletal muscle, or other. Given the strong contribution to depression heritability from the CNS we used LDSC applied to specifically expressed genes (LDSC-SEG) ^110^ across 13 brain regions from the GTEx gene-expression database and brain cell-types (neuronal, astrocyte, oligodendrocyte) using specifically expressed genes derived from mouse-forebrain ^111^.

### Gene and gene-set analyses

A gene-based analysis was applied to the results from our meta-analysis using Multi-marker Analysis of GenoMic Annotation (MAGMA) ^22^ to assess the simultaneous effect of multiple genetic variants on 17,842 genes. To account for LD the European panel of the 1,000 Genomes data (phase 3) ^112^ was used as a reference panel. Genetic variants were assigned to genes based on their position according to the NCBI 37.3 build. To identify those genes that were genome-wide significant a *P*-value threshold was calculated by applying a Bonferroni correction (α = 0.05 / 17,842; *P* < 2.80 × 10^-6^). Regional visualisation plots were produced using the online LocusZoom platform^113^.

A gene-set analysis was then performed on our gene-based results using gene annotation files from the Gene Ontology Consortium (http://geneontology.org/) ^114^ and the Molecular Signatures Database v5.2 ^115^. The annotation file includes gene-sets covering three ontologies; molecular function, cellular components, and biological function and consisted of 5,917 gene-sets. To correct for multiple testing, we used the MAGMA default setting of 10,000 permutations, and applied a Bonferroni correction (α = 0.05 / 5,917; *P* < 8.45 × 10^-6^). Additional gene-sets were obtained from Skene, et al. ^23^ providing expression-weighted enrichment for seven brain cell-types (astrocytes ependymal, endothelial mural, interneurons, microglia, oligodendrocytes, somatosensory pyramidal neurons, and hippocampus CA1 pyramidal neurons). These brain cell-types were assessed in MAGMA using the default setting of 10,000 permutations with a Bonferroni correction (α = 0.05 / 7; *P* < 7.14 × 10^-3^) used to assess significance.

### Drug - gene interactions

We examined genes that were identified as significantly associated (*P* < 2.80 × 10^-6^) with depression using MAGMA ^22^ for interactions with prescribed medications using the Drug Gene Interaction Database (dgidb.genome.wustl.edu) v3.0 ^24^. The Anatomical Therapeutic Chemical (ATC) classification system was used to determine the second level classification of each drug identified. ATC classifications for each drug were obtained from the Kyoto Encyclopaedia of Genes and Genomics (https://www.genome.jp/kegg/drug/) as the primary source of information with additional classifications obtained from the World Health Organisation Collaborating Centre for Drug Statistics Methodology (https://www.whocc.no/atc_ddd_index/). Visualisation of the number of interactions between each gene associated with depression and each second level ATC classification was undertaken using the R package circlize v0.4.1 ^116^.

## Acknowledgements

This research has been conducted using the UK Biobank resource – application number 4844. We are grateful to the UK Biobank and all its voluntary participants. The UK Biobank study was conducted under generic approval from the NHS National Research Ethics Service (approval letter dated 17th June 2011, Ref 11/NW/0382). All participants gave full informed written consent. The authors acknowledge the help, advice, and support from all members of the UK Biobank Psychiatric Genetics Group. The BiDirect cohort and the Münster cohort were approved by the ethics committee of the University of Münster and the Westphalian Chamber of Physicians in Münster, North-Rhine-Westphalia, Germany and written informed consent was obtained from all participants. Ethics approval for the Generation Scotland was given by the NHS Tayside committee on research ethics (reference 15/ES/0040) and all participants provided written informed consent for the use of their data.

We are grateful to the participants and research teams behind the Psychiatric Genomics Consortium, UK Biobank and 23andMe. We thank the following members of the 23andMe Research Team: Michelle Agee, Babak Alipanahi, Adam Auton, Robert K. Bell, Katarzyna Bryc, Sarah L. Elson, Pierre Fontanillas, Nicholas A. Furlotte, Barry Hicks, Karen E. Huber, Ethan M. Jewett, Yunxuan Jiang, Aaron Kleinman, Keng-Han Lin, Nadia K. Litterman, Matthew H. McIntyre, Joanna L. Mountain, Elizabeth S. Noblin, Carrie A.M. Northover, Steven J. Pitts, G. David Poznik, J. Fah Sathirapongsasuti, Olga V. Sazonova, Janie F. Shelton, Suyash Shringarpure, Joyce Y. Tung, Vladimir Vacic, Xin Wang, and Catherine H. Wilson. We would like to thank Dr. Nathan Skene for his advice on analysing the expression-weighted enrichment for brain cell-types and Professor Nick Martin for his suggestions on polygenic risk scores.

AMMcI, and IJD acknowledge support from the Wellcome Trust (Wellcome Trust Strategic Award “STratifying Resilience and Depression Longitudinally” (STRADL) Reference 104036/Z/14/Z and the Dr Mortimer and Theresa Sackler Foundation. IJD is supported by the Centre for Cognitive Ageing and Cognitive Epidemiology, which is funded by the Medical Research Council and the Biotechnology and Biological Sciences Research Council (MR/K026992/1). This investigation represents independent research part-funded by the National Institute for Health Research (NIHR) Biomedical Research Centre at South London and Maudsley NHS Foundation Trust and King’s College London. The views expressed are those of the authors and not necessarily those of the NHS, the NIHR or the Department of Health. GH is supported by the Wellcome Trust [208806/Z/17/Z]. DJS supported by Lister Institute Prize Fellowship 2016-2021. NRW acknowledges NMHRC grants 1078901 and 1087889. The BiDirect Study is supported by grants of the German Ministry of Research and Education (BMBF) to the University of Muenster (01ER0816 and 01ER1506). The Münster cohort was funded by the German Research Foundation (DFG, grant FOR2107 DA1151/5-1 and DA1151/5-2 to UD; SFB-TRR58, Projects C09 and Z02 to UD) and the Interdisciplinary Center for Clinical Research (IZKF) of the medical faculty of Münster (grant Dan3/012/17 to UD). The PGC has received major funding from the US National Institute of Mental Health and the US National Institute of Drug Abuse (U01 MH109528 and U01 MH1095320).

## Author Contributions

D.M.H., D.J.S., G.B., C.M.L., and A.M.McI. conceived the research project. M.J.A., J.R.I.C., J.W., D.J.S., G.B., and A.M.McI. determined the variables that formed the depression phenotypes within UK Biobank. D.M.H., M.J.A., J.R.I.C., R.E.M., and G.D. applied quality control to the UK Biobank data. M.J.A. ran the association analysis in UK Biobank. 23andMe R.T. provided the summary statistics from the Hyde, et al. ^8^ analysis. M.D.D.W.G.P.G.C., M.T., S.R., P.F.S., N.R.W., and C.M.L. provided the summary statistics from the Wray, et al. ^9^ analysis with additional removal of overlapping cohorts. D.M.H. ran the meta-analysis of the three contributing cohorts. C.T. and D.A.H. conducted the replication analysis in 23andMe for the significant variants in the meta-analysis. TK.C., S.P.H and E.Y.X. conducted the polygenic risk score analysis. K.B., H.T. and R.R. ascertained the BiDirect cohort and the Münster cohort was ascertained by V.A., B.T.B., U.D., and K.D. T-K.C. and D.M.H calculated the genetic correlations and partitioned the heritability component of depression. D.M.H. and G.H. performed the Mendelian randomization analyses. D.M.H. ran the MAGMA-based analyses with J.W., M.S., J.G., E.M.W., C.A., X.S., and M.C.B. examining the genes and gene-sets identified. J.D.H., D.M.H., and A.M.McI. conducted the gene x drug interaction analysis. I.J.D., D.J.P., H.C.W., S.R., D.J.S., P.F.S., N.R.W., E.M.B., G.B., C.M.L., and A.M.McI. provided expertise of association study methodology and statistical analysis. D.M.H. oversaw the research project and serves as the primary contact for all communication. All authors commented on the manuscript.

## Data availability

Summary statistics for 10,000 genetic variants from the meta-analysis of 23andMe_307k, UK Biobank, and PGC_139k and the summary statistics for all assessed genetic variants for the meta-analysis of depression in UK Biobank and PGC_139k are available from: http://dx.doi.org/10.7488/ds/2458.

To access the summary statistics for all assessed genetic variants for the meta-analysis of depression in 23andMe_307k, UK Biobank, and PGC_139k a data transfer agreement is required from 23andMe before a request should be made to the corresponding author.

The raw genetic and phenotypic UK Biobank data used in this study, which were used under license, are available from: http://www.ukbiobank.ac.uk/.

The genome-wide summary statistics for the Hyde et al. analysis of 23andMe, Inc. data were obtained under a data transfer agreement. Further information about obtaining access to the 23and Me, Inc. summary statistics are available from: https://research.23andme.com/collaborate/

The genome-wide summary statistics for the Wray et al. analysis of PGC data were obtained under secondary analysis proposals #60 and #76. Further information about obtaining access to the PGC summary statistics are available from: http://www.med.unc.edu/pgc/statgen

## Supplementary Information

General directionality agreement of variants that have previously been associated with depression

Cohort information for the 23andMe replication dataset

Members of the Major Depressive Disorder Working Group of the Psychiatric Genomics Consortium and their affiliations

Supplementary Figure 1

Quantile-quantile plot of the observed *P*-values on those expected for each genetic variants association with depression in our meta-analysis

Supplementary Figure 2

Odds ratios and 95% confidence intervals for Major Depressive Disorder (MDD) in Generation Scotland based on polygenic risk score (PRS) deciles calculated from the current meta-analysis of depression and from the summary statistics from the genome-wide association study of major depression conducted by Wray, et al. ^9^.

Supplementary Figure 3

Significant genetic correlations (r_G_; *P* < 0.01, after false discovery rate correction) between depression and other behavioural and disease related traits using LD score regression implemented in LD Hub software (http://ldsc.broadinstitute.org/).

A negative r_G_ indicates that an earlier or lower value of a continuous trait (i.e. earlier father’s age of death or lower subjective well being was associated with depression. A positive r_G_ indicates that a later or higher value of a continuous trait (i.e. higher triglyceride level) was associated with depression. Where multiple studies have examined a single trait the pubmed number of the study is given in brackets.

Supplementary Figure 4

Mendelian randomisation test for a putative causal effect of depression on neuroticism using inverse weighted regression, MR Egger and a weighted median test

Supplementary Figure 5

Mendelian randomisation analysis for a putative causal effect of depression on ever vs. never smoked using inverse weighted regression

Supplementary Figure 6

Mendelian randomisation test for a putative causal effect of neuroticism on depression using inverse weighted regression, MR Egger and a weighted median test

Supplementary Figure 7

Contribution of functional annotation categories to the heritability of depression based on the variants within each category

Error bars represent jackknife standard errors for each the estimate of enrichment, and an asterisk indicates significance (*P* < 0.0021) after Bonferroni correction. The dashed line represents the threshold for no enrichment.

Supplementary Figure 8

Coefficients (β) of significantly enriched brain cell regions using GTEx overlaid on physical representation of brain anatomy

The pseudo-coloring highlights the coefficients of the brain regions in red that were significantly enriched (*P* < 0.05) for depression variants

Supplementary Figure 9

Stratified LD score regression analyses showing significance of enrichment estimates for 3 brain cell types in depression.

The dashed line represents the Bonferroni threshold for significance (*P* < 0.0167) and * indicates significant enrichment for that brain cell type

Supplementary Figure 10

Regional visualization plot centred on the independently-associated variant, rs1021363, close to the Sortilin related VPS10 domain containing receptor 3 (*SORCS3*) gene on chromosome 10. Recombination rates used in the plots are based on the European 1000 Genomes panel from November 2014

Supplementary Figure 11

Regional visualization plots centred on independently-associated variants (A. rs2568958 and B. rs10890020) close to the Neuronal Growth Regulator 1 (*NEGR1*) gene on chromosome 1. Recombination rates used in the plots are based on the European 1000 Genomes panel from November 2014

Supplementary Figure 12

Regional visualization plots centred on independently-associated variants (A. rs62091461, B. rs12966052, and C. rs12967143) close to the Transcription Factor 4 (*TCF4*) and *RAB27B* genes on chromosome 18. Recombination rates used in the plots are based on the European 1000 Genomes panel from November 2014

Supplementary Table 1

Variants with a *P*-value < 5 × 10^-8^ for an association with depression in the meta-analysis of PGC_139k, 23andMe_307k and UK Biobank

Supplementary Table 2

The direction of effect of previously reported significant variants for depression across the studies contributing to the meta-analysis

Supplementary Table 3

Genetic correlations between depression and other behavioural and disease related traits using LD score regression implemented in LD Hub software (http://ldsc.broadinstitute.org/)

Supplementary Table 4

Mendelian randomization analysis between depression and other traits using MR Egger test for directional horizontal pleiotropy, inverse variance weighted (IVW) test for variant heterogeneity and IVW regression, weighted median and MR Egger tests for a causal effect

Supplementary Table 5

Heritability partitioned by functional annotation enrichment. The asterisk indicates significance after Bonferroni correction for multiple testing (*P* < 0.0021)

Supplementary Table 6

Partitioning of the heritability estimate by cell type enrichment. The asterisk indicates significance after Bonferroni correction for multiple testing (*P* < 0.0056)

Supplementary Table 7

Enrichment estimates for brain regions using GTEx. The asterisk indicates significance after Bonferroni correction for multiple testing (*P* < 0.0038)

Supplementary Table 8

Enrichment estimates for brain cell types. The asterisk indicates significance after Bonferroni correction for multiple testing (*P* < 0.0167)

Supplementary Table 9

Genome-wide significant gene-based hits (*P* < 2.80 x 10-6) in the meta-analysis of depression using MAGMA. NSNPS is the number of SNPs in the gene; NiSNPs is the number of independent SNPs in the gene

Supplementary Table 10

Number and proportion of gene overlap within the Gene Ontology Consortium gene-sets associated (*P*_corrected_ < 0.05) with depression

Values on the lower diagonal are the number of overlapping genes between gene sets. Values on the upper diagonal are the proportion of overlapping genes within the gene set containing the lower number of genes

Supplementary Table 11

Drug x gene interactions for the genes identified as significantly associated with depression with interactions obtained from the drug gene interaction database (http://dgidb.genome.wustl.edu/). The Anatomical Therapeutic Chemical (ATC) classification for each drug is provided along with the type of interaction and its source

